# *Ex-vivo* mouse precision cut tumour slices for modelling hepatocellular carcinoma; A 3Rs solution for at-scale drug screening

**DOI:** 10.1101/2024.09.16.613213

**Authors:** Amy L Collins, Keara Kirkness, Erik Ramon-Gil, Eleni Tzortzopoulou, Daniel Geh, Rainie Cameron, Saimir Luli, Eman Khurram, Daniel Storey, Hannah Paish, David McDonald, Andrew Filby, Lee A Borthwick, Fiona Oakley, Derek Mann, Jack Leslie

## Abstract

Disease modelling is vital for improving knowledge of disease mechanisms and for development of new therapeutic molecules and strategies. Modelling the intact living tumour microenvironment (TME) is increasingly considered to be vital not only for gaining a better understanding of the biology of cancer but for examining the efficacy of novel oncology drugs. To date, pre-clinical mouse models of cancer have represented the mainstay methodology for studying the evolving TME and for determining the effects of potential therapeutic molecules on tumour evolution and growth. Regarding drug screening, *in vivo* mouse models are expensive, require the use of large cohorts of mice and involve the administration of drugs with unknown toxicities to animals which often result in adverse effects that can cause animal suffering and the discontinuation of drug investigations. Hepatocellular carcinoma (HCC) is a primary cancer of the liver for which there is an urgent need for improved systemic treatments due to the disease usually being diagnosed at an advanced stage and current treatments having limited efficacy. To provide a practical solution to the screening of drugs for their likely efficacy in HCC we have developed an *ex-vivo* model in which orthotopic tumours are excised from the liver and subsequently processed to generate precision-cut tumour slices (PCTS) which provide an intact culture model of the HCC-TME. We describe simplified culture conditions that maintain the viability and metabolic activity of live PCTS which maintain the architecture, cellular complexity, drug sensitivity and responsiveness to immunotherapy of the original tumour. Importantly, we show that HCC derived PCTS can be miniaturised to 96-well scale and modified to express soluble luciferase, which in combination enabled non-destructive screening of a library of 26 drugs at two doses using just 5 tumours as the source for PCTS. This screen identified two small molecules, salinomycin and rottlerin, that have potent anti-tumour activities in HCC-PCTS and subsequently validated salinomycin as effective *in vivo*. In summary, we report a 3Rs (reduction, refinement and replacement) solution for study of HCC biology and for 96-well-scale screening of potential therapeutic agents in the context of an intact, metabolically active TME.

## Introduction

Liver cancer presents a growing global health challenge and is one of the few cancer types increasing in incidence. There were approximately 906,000 new cases of liver cancer in 2020, and hepatocellular carcinoma (HCC) comprises roughly 80% of primary liver cancer cases (1, 2). A number of risk factors are associated with the development of HCC, including viral infection with hepatitis B and hepatitis C, metabolic dysfunction-associated steatotic liver disease (MASLD), metabolic associated steatohepatitis (MASH) and alcohol-related liver disease (ALD) (3, 4). The majority of HCC cases can be attributed to chronic liver disease, and cirrhosis is the strongest risk factor for HCC development (5, 6).

Despite the emergence of various therapeutic options for the treatment of advanced HCC over the past two decades, their efficacy is limited and only exert favourable effects in a minority of patients. The combination therapy of atezolizamab (anti-PD-L1) and bevacizumab (anti-VEGF) was demonstrated to be superior to the previous gold standard therapy for advanced HCC, the multi-target tyrosine kinase inhibitor sorafenib (7). However, the survival benefit offered by this combination therapy is still minimal, and just 30% of patients exhibit a therapeutic response. Superior therapeutic strategies are required to target highly aggressive tumours where currently available therapies have limited efficacy in order to attenuate the increasing mortality rates for HCC (8). The tumour immune microenvironment (TME) is a complex network whereby crosstalk between various cellular components contributes to HCC development and therapeutic response (9). Animal models are therefore considered to be an effective method of determining pre-clinical therapeutic efficacy in which components of the tumour microenvironment are accounted for. Orthotopic xenograft mouse models of HCC generate fast-growing tumours suitable for the screening of novel therapies, and it has previously been demonstrated that Hep-53.4 cells generate fibrotic tumours with an abundance of infiltrating T cells and dendritic cells, and are responsive to anti-PD1 immunotherapy (10, 11). Furthermore, transcriptomic analysis identified that Hep-53.4 orthotopic tumours are both poorly differentiated and highly proliferative, indicating that the model may offer utility for therapeutic screening for aggressive HCC tumours (12).

Precision cut liver slices and precision cut tumour slices (PCLS and PCTS respectively) retain the architecture, ECM composition and complex cell/cell interactions of native tissue and provide a platform for studying disease mechanisms and therapeutic response *ex vivo* (13–15). Routinely acquiring large quantities of human tumour tissue for PCTS generation can present difficulties, given the importance of resected tumour samples which must be carefully examined by pathologists so not to compromise patient care. Additionally, the degree of structural and cellular heterogeneity between different patient tumour samples, and even within any one individual HCC tumour, is considerable and therefore makes standardisation of the platform challenging, this being particularly problematic for oncology drug screening and development. As a result, we sought to generate PCTS from orthotopic murine Hep-53.4 tumours to provide a robust, reproducible *ex vivo* model of HCC capable of assessing novel therapeutic strategies to target poorly differentiated HCC. The ability to screen multiple therapeutic combinations in PCTS from the same tumour provides a cost-effective method of assessing drug efficacy in line with the 3R’s: reducing, replacing and refining the use of animals in research.

## Methods

### Mice

All animal experiments were approved by the Newcastle Ethical Review Committee and performed under a UK Home Office licence, in accordance with the ARRIVE guidelines. All mice were housed in the Comparative Biology Centre at Newcastle University in pathogen-free conditions with free access to food and water. C57BL/6 WT mice were purchased from Envigo for *in vivo* and *ex vivo* experiments.

### Orthotopic liver cancer model

Surgeries were performed under general anaesthesia with isoflurane and all animals were administered with pre-and post-surgery buprenorphine (0.003 mg/ml). Following a laparotomy, 1×10^6^ Hep-53.4, Hepa1-6 or H22 HCC cells were implanted into the left lobe of C57BL/6 mice via intrahepatic injection. In vivo imaging systems (IVIS) imaging was performed weekly to assess tumour growth via bioluminescence. After the Hep-53.4 cell line was selected for orthotopic model generation, mice with Hep-53.4 tumours were harvested at days 14, 21 and 28 for characterisation of the tumour and liver tissue. Therapeutic intervention with 45 mg/kg sorafenib (Tocris), 10 mg/kg lenvatinib (Selleckchem) or a PEG/DMSO control was started 14 days post implantation and was administered via daily oral gavage until mice were harvested at day 28. Therapeutic intervention with 4 mg/kg salinomycin (MedChemExpress) or a corn oil/DMSO control was started 14 days post implantation was administered via daily intraperitoneal injection until mice were harvested at day 28.

### Precision Cut Liver and Tumour Slice generation

Liver and tumour tissue was cored using a 3 mm or 8 mm Stiefel biopsy punch (Medisave, Weymouth, UK). Tissue cores were submerged in 3% low gelling temperature agarose (Sigma-Aldrich) in metal moulds and placed at 4°C for 10 minutes to set. Agarose-embedded tissue cores were then cut using a Leica VT1200S vibrating blade microtome (Leica) at a depth of 250 μm. PCLS generated were cultured in 8-µm-pore Transwell inserts in our Bioreactor as previously described (14), or alternatively in standard static 12-well or 96-well culture plates. Tissue was cultured in Williams’ Medium E (Sigma-Aldrich) supplemented with 1% penicillin-streptomycin, 1% L-glutamine, 1% pyruvate, 1 x insulin transferrin-selenium X, 2% fetal bovine serum and 100 nM dexamethasone, at 37°C supplemented with 5% CO_2_. Media was completely replaced daily.

### Tissue culture treatments

All PCTS and PCLS were cultured in supplemented Williams’ Medium E alone for 1 day to allow the tissue to stabilise. To investigate TKI response, PCTS and PCLS 8 mm in diameter were cultured in BioR plates in the Bioreactor and treated with sorafenib (10 µM – 20 µM) (Tocris), lenvatinib (0.25 µM – 1.0 µM) (Selleckchem) or DMSO control media for a further 3 days after the rest period. To model immunotherapy responses, PCTS 8 mm in diameter were cultured in BioR plates in the Bioreactor, and treated with 20 µg/ml Ultra-LEAF™ Purified anti-mouse CD279 (PD-1) (Biolegend) or rat IgG2a, κ isotype control (Biolegend) from day 2 until the PCTS were harvested at day 4. To screen a panel of 26 compounds (**Figure 6A**), PCTS 3 mm in diameter were cultured in static 96-well culture plates and treated with two doses of each compound or controls for a further 7 days after the rest period. To investigate salinomycin toxicity PCLS and PCTS 3 mm in diameter were cultured in BioR plates in the Bioreactor with DMSO control, salinomycin (5 µM – 10 µM) or sorafenib (20 µM) for a further 3 days (PCLS) or 7 days (PCTS) after the rest period.

### Cell culture and transfection

Hep-53.4 and Hepa1-6 cells were cultured in DMEM with high glucose (Sigma), and H22 cells were cultured in RPMI-1640 (Sigma), at 37°C with 5% CO_2_. All culture media was supplemented with 10% FBS, 1% penicillin-streptomycin, 1% L-glutamine and 1% pyruvate. Cells were routinely screened for mycoplasma and were mycoplasma negative.

Stable transfections were performed Hep-53.4 cells using a custom pNL(sNLuc/CMV/NeoR) (**Supplemental Figure 1**) expression vector using the Lipofectamine 3000 Transfection Reagent kit (Thermo Fisher). 100,000 cells were seeded in each well of a 6-well plate. The following day, a transfection complex comprised of 250 µl Opti-MEM™ (Thermo Fisher), 5 µl P3000 Reagent, 7.5 µl Lipofectamine 3000 Reagent and 2.5 µg plasmid DNA was mixed and left for 10 minutes to enable DNA-lipid formation. The cell culture media was refreshed, and the transfection complex was added to the cells in a drop-wise manner. The following day, the media was removed from all cells and replaced with selection media containing 1 mg/ml G418 disulfate salt (Sigma). The cells remained in selection media until all cells in a control well had visibly died. Transfected cell lines were cultured in selection media approximately once every two months to prevent an expansion of WT cells.

### Colorimetric assays

CyQUANT™ LDH Cytotoxicity Assay (Thermo Fisher) was performed as per the manufacturers’ instructions.

### Enzyme linked immunosorbent assay (ELISA)

Sandwich ELISA quantification for mouse albumin (ab210890, Abcam) was performed according to the manufacturers’ instructions on culture media samples harvested daily from PCLS.

### Resazurin assay

4.5 mM resazurin stock solution was diluted in Williams’ Medium E to a working concentration of 450 µM. Each tissue slice was incubated in a 96-well plate with 100 µl of the working resazurin solution for 1 hour at 37°C with 5% CO2. 100 µl of the working resazurin solution was used as a negative control. After 1 hour, the solution was transferred to an opaque 96-well plate (Greiner) and the fluorescence was measured on a Filtermax F5 multi-mode plate reader, ex 535 nm and em 595 nm.

### Luminescent assays

Levels of secreted luciferase in the PCTS culture media from SecLuc transfected HCC cells, was determined using the Nano-Glo® Luciferase Assay System (N1120, Promega). Nano-Glo® Luciferase Assay Reagent was prepared as per the manufacturers’ instructions. 40 µl of undiluted cell culture media samples were added to an opaque 96-well plate (Greiner) and 40 µl of the relevant culture media was used as a negative control. The samples were combined in a 1:1 ratio with the Nano-Glo® Luciferase Assay Reagent and left for at least 3 minutes for the reaction to take place. The luminescence values were then measured using Filtermax F5 multi-mode plate reader. Cell Titer-Glo® 3D Cell Viability Assay (Promega) was performed as per the manufacturers’ instructions.

### RNA isolation, cDNA synthesis and PCR

The QIAGEN RNeasy Mini Kit (74106; QIAGEN) was used to extract RNA from tissue as per the manufacturers’ instructions. 1 µg RNA was treated with DNase (Promega) and cDNA was synthesised via incubation with random hexamer primer (C1181; Promega) and M-MLV reverse transcriptase (M1701; Promega). Real-time PCR was carried out using SYBR Green JumpStart Taq ReadyMix (S4438; Sigma) as per the manufacturers’ instructions with primers previously described (16).

### Histology and immunohistochemistry

Frozen 10 µm thick tissue sections were stained with Oil Red O, and formalin-fixed, paraffin-embedded 5 µm thick tissue sections were stained with haematoxylin and eosin using established protocols. To perform immunohistochemistry, sections were first deparaffinised and endogenous peroxidase activity was blocked with 0.6% hydrogen peroxide/methanol solution. Antigen retrieval was performed using either antigen unmasking solution (Vector Laboratories) for αSMA (1:1000; F3777; Sigma-Aldrich), Active Caspase-3 (1:400; 9661; Cell Signaling), Cytokeratin 19 (1:250; ab85632; Abcam), Ly6G (1:200, ab210204; Abcam) and PCNA (1:3000; ab18197; Abcam), or 1 mM Tris-EDTA (pH 9.0) for CD3 (1:200; MCA1477; Bio-Rad), CD79A (1:5000; ab300150; Abcam), Cytokeratin 18 (1:800; ab181597; Abcam), F4/80 (1:800; 70076; Cell Signaling), Ki-67 (1:10,000; 14-5698-82; Thermo Fisher) or VCAM-1 (1:200; 32653S; Cell Signaling). The Avidin/Biotin Blocking Kit (Vector Laboratories) was used to block endogenous avidin and biotin, followed by blocking of non-specific binding with Normal Goat Serum (Vector Laboratories) for 45 minutes. Tissue sections were then incubated with primary antibodies at the relevant dilutions overnight at 4°C. The slides were washed and sections were incubated with biotinylated goat anti-rabbit (1:600; Vector Laboratories), biotinylated goat anti-rat (1:200; Serotec) or biotinylated goat anti-fluorescein (1:300; Vector Laboratories) secondary antibodies for 45 minutes. Slides were washed and incubated with Vectastain Elite ABC HRP Reagent (Vector Laboratories) for 30 minutes. Staining was developed using DAB peroxidase substrate kit (Vector Laboratories), followed by counterstaining with Mayer’s haematoxylin. Sections were then dehydrated and mounted in Pertex Mounting Medium (Cell Path). Terminal deoxynucleotidyl transferase-mediated dUTP nick end (TUNEL) labelling was performed using the TUNEL Assay Kit – HRP-DAB (Abcam) as per the manufacturer’s instructions. Tissue sections were analysed at 20x magnification using a Nikon Eclipse Ni-U microscope and NIS-Elements BR analysis software. A minimum of 12 non-overlapping fields were analysed from in vivo tissue sections, and a minimum of 6 non-overlapping fields were analysed from PCLS and PCTS due to the smaller available area of tissue.

### Hyperion imaging mass cytometry

Hyperion Imaging Mass Cytometry (IMC) was performed on formalin-fixed paraffin-embedded (FFPE) 5 µm thick tissue sections. Slides were first deparaffinised and rehydrated by passing the sections through clearene, followed by 100%, 90%, 70% and 50% ethanol solution for 5 minutes each. Sections were then washed in deionised water for 5 minutes, before heat-mediated antigen retrieval was performed using Tris-EDTA solution at pH 9.0. The sections were allowed to cool and then washed in PBS for 5 minutes. A ring was drawn around each tissue section with a hydrophobic pen and non-specific binding was blocked with 3% BSA/PBS for 45 minutes. 200 µl of a metal-conjugated CD8 (rabbit monoclonal, clone EPR21769, Abcam) and Ki67 (Rat monoclonal, clone SolA15, ThermoFisher) primary antibodies was then added to each section in 0.5% BSA/PBS and incubated overnight at 4°C. CD8 and Ki67 primary antibodies were labelled with 164 Dy and 163 Dy metal isotopes respectively using the Maxpar® X8 Multimetal Labeling Kit—40 Rxn (201300, Standard Biotools). The sections were washed in Tris-Buffered Saline + 0.1% Tween (TBS-T) followed by two consecutive washes in PBS. The sections were then incubated for 30 minutes with 125 µM (193lr) Intercalator at a dilution of 1:400. The sections were washed in ultra-pure water for 5 minutes and subsequently air-dried at room temperature. A region of interest (ROI) was selected around the HCC spheroid and the tissue was ablated by the Hyperion Fluidigm Imaging System.

## Results

### Characterisation of an orthotopic HCC model for PCTS production and pre-clinical drug testing

Given the challenges in acquiring large quantities of fresh patient HCC tumour resection material suitable for generating PCTS, we first developed a rapid syngeneic orthotopic HCC model which was suitable for makings PCTS. To establish an orthotopic model, we identified three commercially available murine cell lines of hepatocellular origin, luciferase tagged them and intrahepatically injected 1×10^6^ cells into the large lobe of the liver following a laparotomy under surgical anaesthesia **(Figure 1A)**. Tumour engraftment and growth was then monitored weekly by bioluminescence IVIS imaging. Only Hep-53.4 cells resulted in 100% engraftment, an exponential increase in bioluminescence signal and the reproducible formation of large macroscopic tumours **(Figure 1B-E)**.

**Figure 1.**
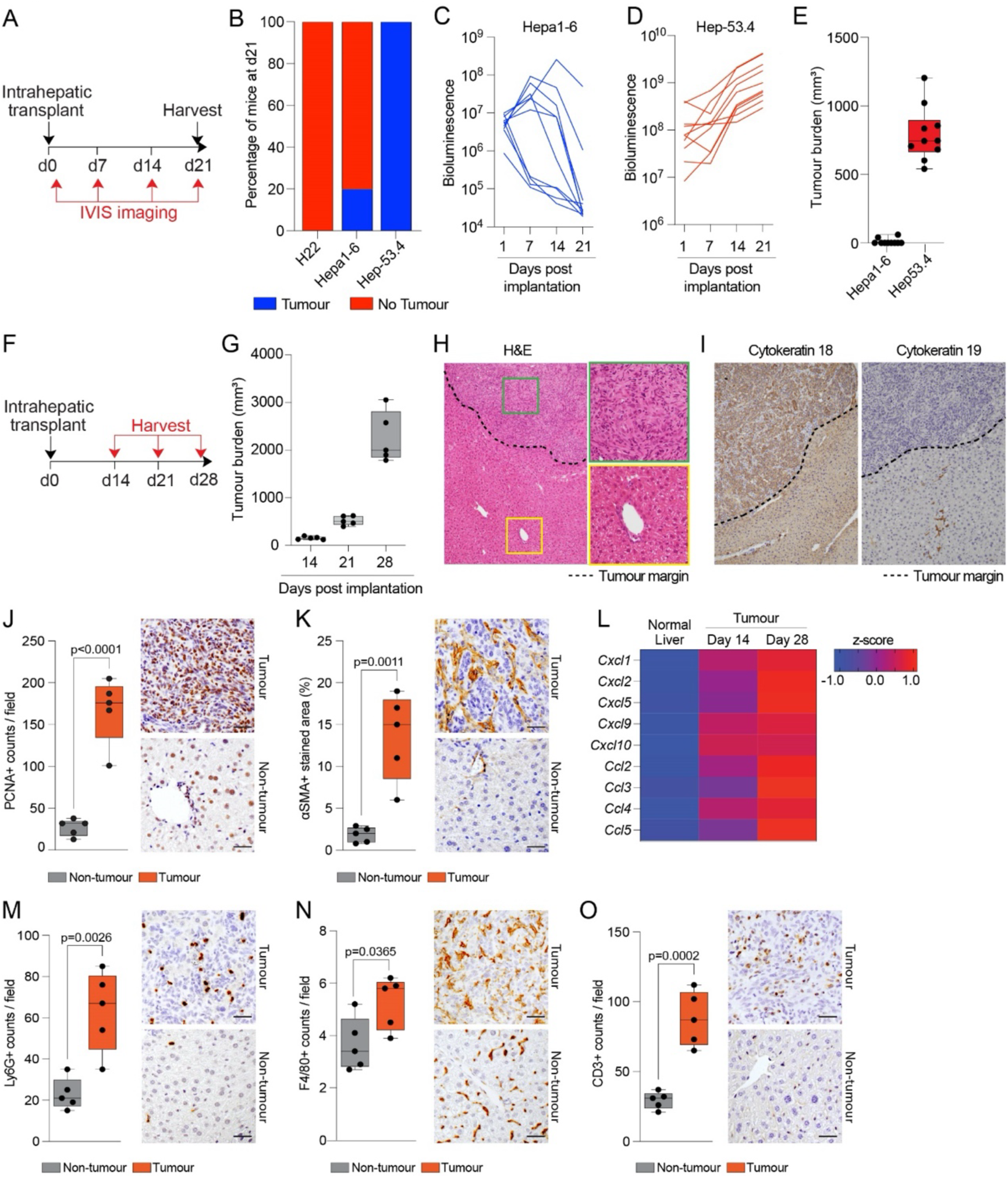
Hep-53.4 cells generate large, fast-growing orthotopic tumours. (A) Schematic shows the timeline of intrahepatic transplant of Hep-53.4, Hepa1-6 or H22 cells and subsequent weekly IVIS imaging to assess HCC engraftment and growth. (B) Percentage stacked bar chart shows the percentage of mice that developed tumours 21 days after intrahepatic transplant of Hep-53.4, Hepa1-6 or H22 cells. N=10 mice per cell line. (C-D) Graphs show bioluminescence levels from IVIS imaging of mice at 1, 7, 14 and 21 days following intrahepatic injection of (C) Hepa1-6 or (D) Hep-53.4 cells. (E) Graph shows tumour burden of Hepa1-6 and Hep-53.4 orthotopic tumours 21 days after intrahepatic transplant. Data are mean ± s.e.m. from N=10 mice per cell line. (F) Schematic shows the timeline of intrahepatic transplant of Hep-53.4 cells and subsequent harvest at days 14, 21 and 28. (G) Graph shows tumour burden of Hep-53.4 tumours harvested 14, 21 and 28 days after intrahepatic transplant. Data are mean ± s.e.m. from N=5 mice per time point. (H-I) Representative images of (H) H&E-stained and (I) cytokeratin-18- and cytokeratin-19-stained tumour and non-tumour tissue. Black dotted line denotes tumour margin. (J-K) Histological quantification and representative images of (J) PCNA-stained and (K) αSMA-stained tumour and non-tumour tissue harvested 28 days after intrahepatic transplant. Data are mean ± s.e.m. from N=5 mice. (L) Heatmap showing expression of *Cxcl1, Cxcl2, Cxcl5, Cxcl9, Cxcl10, Ccl2, Ccl3, Ccl5* and *Ccl5* in normal liver tissue and Hep-53.4 tumour tissue 14 and 28 days after intrahepatic transplant. (M-O) Histological quantification and representative images of (M) Ly6G-stained, (N) F4/80-stained and (O) CD3-stained tumour and non-tumour tissue harvested 28 days after intrahepatic transplant. Data are mean ± s.e.m. from n = 5 mice. Scale bars: 50 μm.

We next determined the kinetics and humane endpoints of macroscopic tumour development in the Hep-53.4 model. Mice received a single intrahepatic injection of 1×10^6^ Hep-53.4 cells and were harvested weekly **(Figure 1F)**. Small macroscopic tumours were observed at 14 days post injection **(Figure 1G)**. Tumour growth was exponential, increasing significantly between day 21 and day 28, after which animals typically needed to be humanely killed due to tumour burden **(Figure 1G)**. Histological analysis of the Hep-53.4 tumours revealed localisation of the tumour primarily to one half of the left lobe, with thickening of the hepatic plate visible (**Figure 1H**). Immunohistochemical staining for cytokeratin 18 and cytokeratin 19 revealed that Hep-53.4 tumours are of hepatocellular origin and are not positive for the biliary marker cytokeratin 19 (**Figure 1I**). Immunohistochemical staining for PCNA^+^ proliferating cells and αSMA^+^ stromal cells revealed that Hep-53.4 tumours are highly proliferative and display significant cancer associated fibroblast activation **(Figure 1J-K)**.

We have previously shown by whole exome sequencing that Hep-53.4 cells are mutant for p53, WT for CTNNB1 as well as APC and AXIN2 suggesting that these tumours were likely to be immunogenic and would potentially respond to immunotherapy (17). Analysis of Hep-53.4 tumours revealed they were highly immunogenic displaying increased proinflammatory gene expression **(Figure 1L)** and increased numbers of infiltrating Ly6G^+^ neutrophils, F4/80^+^ macrophages and CD3^+^ T cells compared to adjacent normal liver **(Figure 1M-O)**.

### PCTS retain characteristics of *in vivo* tumours in culture

PCLS are an *ex vivo* culture system that have been used to study hepatic fibrosis and cancer as well as drug metabolism (14, 15, 18). We therefore generated PCLS and PCTS from our highly characterised, aggressive HCC mouse model and cultured them in our patented bioreactor system, as previously described (**Figure 2A**)(14). Like human and rat PCLS, mouse PCLS and PCTS cultured in the bioreactor system were metabolically active following 4 days in culture (**Figure 2B**). Analysis of LDH, indicative of necrosis, showed that cell death caused as a result of tissue slicing began to stabilise after 24 hours in culture, and of note we observed lower levels of cell death in PCTS compared with PCLS (**Figure 2C**). Tissue morphology and integrity, evaluated by H&E staining, was maintained up to day 4 in Bioreactor cultured PCLS and PCTS (**Supplemental Figure 2A**). However, mouse PCLS viability was observed to begin to deteriorate after 3 days in culture (**Supplemental Figure 2B and Supplemental Figure 4D**), possibly due to the fragility of the tissue, this contrasting with stable viability for up to 6 days for rat and human PCLS maintained in the rocked bioreactor (14). However, PCTS remained metabolically active for at least 10 days with no obvious necrosis despite the increased cellular density present in PCTS compared to PCLS (**Figure 2D**). PCTS cultured for 4 days maintained an elevated number of PCNA^+^ tumour cells, a dense αSMA^+^ stroma and increased expression of cell cycle genes compared to non-tumour PCLS (**Figure 2E-G**). The tumour slices maintained both CD3^+^ T cell and F4/80^+^ macrophages when cultured for 4 days, albeit with a modest reduction in their numbers being observed after 2 days **(Figure 2H-I)**. By contrast Ly6G^+^ neutrophils were only present in the tissue for only the first 24 hours of culture due to their short lifespan **(Figure 2J)**. Taken together these observations suggest that bioreactor cultured PCTS represent a potential *ex vivo* platform for studying an intact metabolically active tumour microenvironment.

**Figure 2.**
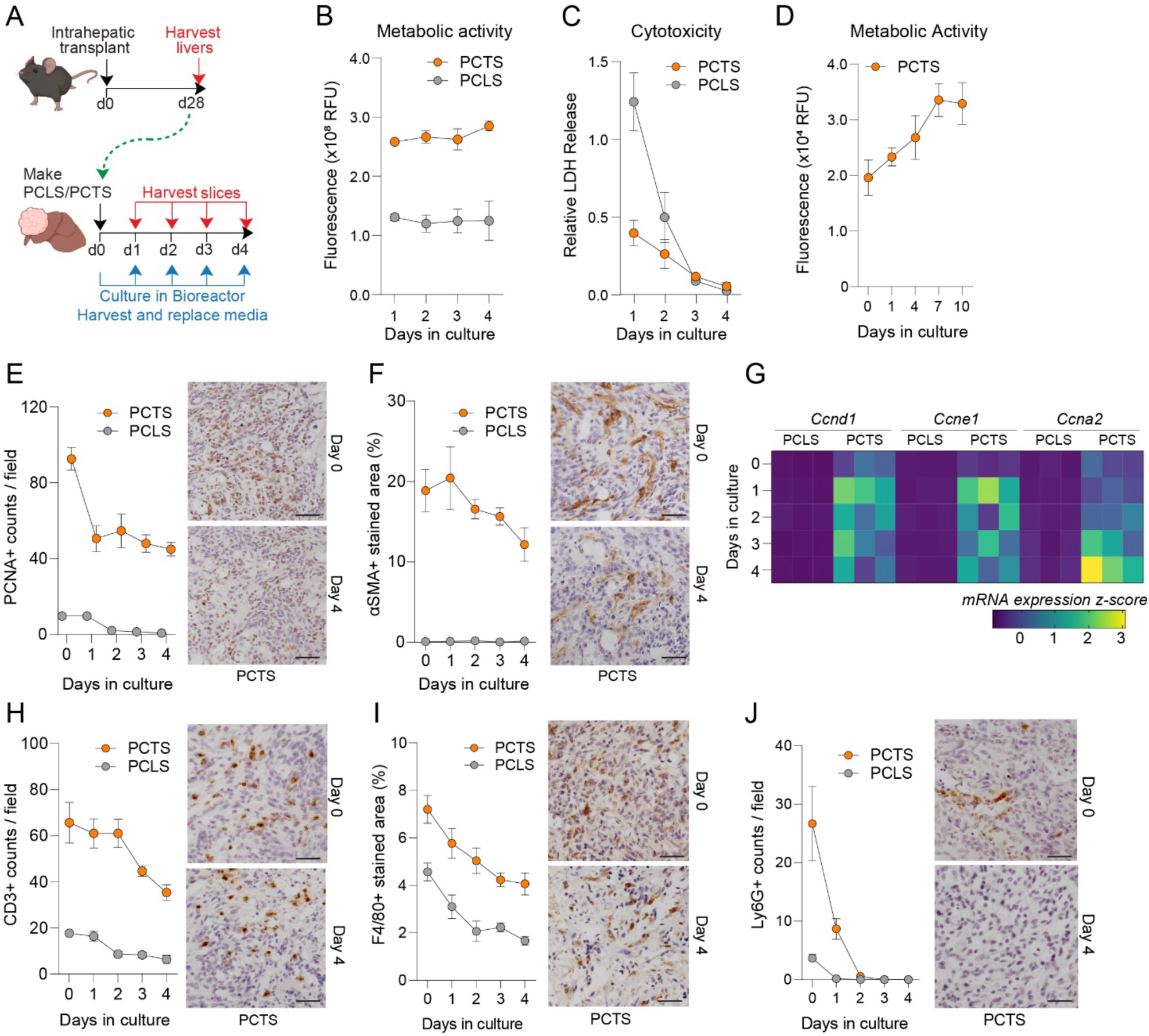
PCTS retain characteristics of original tumour in culture. (A) Schematic shows the timeline of intrahepatic transplant of Hep-53.4 cells to generate orthotopic tumours for generation of PCTS and PCLS, and subsequent 4-day culture period in a rocked Bioreactor platform. (B-C) Graphs show levels of (B) metabolic activity measured via resazurin assay and (C) cytotoxicity measured via LDH assay from PCTS and PCLS cultured for 4 days in a rocked Bioreactor platform. Data are mean ± s.e.m. from N=3 PCTS/PCLS. (D) Graph shows metabolic activity measured via resazurin assay of PCTS cultured for 10 days in a rocked Bioreactor platform. Data are mean ± s.e.m. from up to N=6 PCTS per time point. (E-F) Histological quantification and representative images of (E) PCNA-stained and (F) αSMA-stained PCTS and PCLS cultured for 4 days in a rocked Bioreactor platform. Data are mean ± s.e.m. for N=3 PCTS/PCLS per time point. (G) Heatmap showing expression of cell cycle genes *Ccnd1, Ccne1* and *Ccna2* in PCLS and PCTS between 0 and 4 days in culture. (H-J) Histological quantification and representative images of (H) CD3-stained, (I) F4/80-stained and (J) Ly6G-stained PCTS and PCLS cultured for 4 days in a rocked Bioreactor platform. Data are mean ± s.e.m. from N=3 PCTS/PCLS per time point. Scale bars: 50 μm.

### PCTS recapitulate *in vivo* responses to HCC systemic therapies

We next asked if orthotopic HCC responds to the tyrosine kinase inhibitors (TKIs) sorafenib and lenvatinib, which are both first-line treatments for advanced HCC (**Figure 3A**). Sorafenib targets serine/threonine and tyrosine kinases in multiple oncogenic and angiogenic signalling pathways, resulting in its well characterised cyto-static, apoptotic and anti-angiogenic effects (19, 20). Lenvatinib on the other hand is primarily thought to enact its anti-tumour effects by suppressing angiogenesis via targeting VEGFR1-3, although cyto-static effects have been described *in vitro* (21). Compared to vehicle control, both sorafenib and lenvatinib treated mice displayed a significant reduction in tumour burden and liver weight (**Figure 3B and Supplemental Figure 3A**). This therapeutic effect was associated with a significant reduction in proliferating Ki-67^+^ cells in lenvatinib treated mice and a non-significant increase in active caspase-3^+^ apoptotic tumour cells in sorafenib treated mice (**Figure 3C-D).**

**Figure 3.**
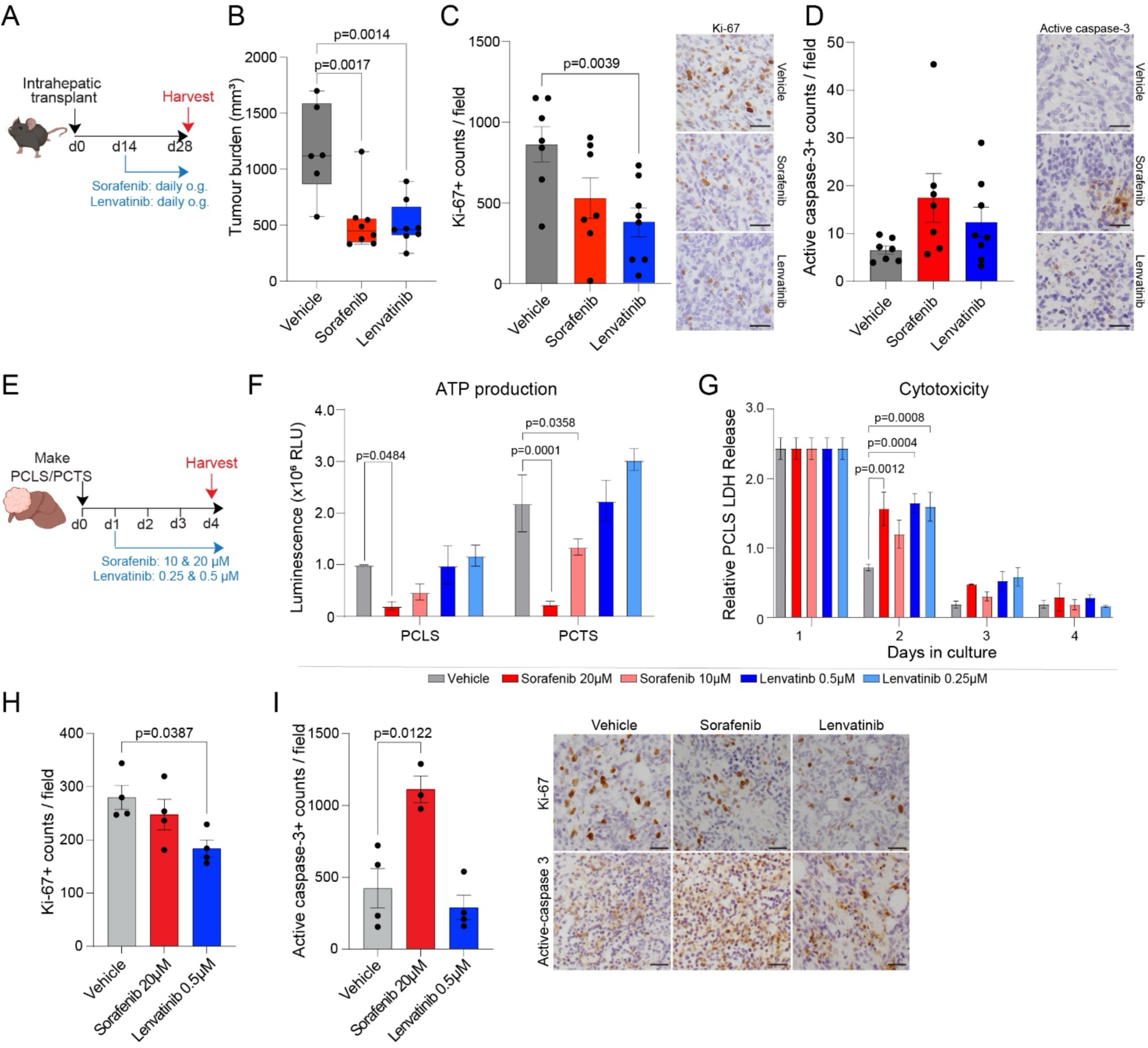
PCTS recapitulate in vivo responses to HCC tyrosine kinase inhibitors. (A) Schematic shows the timeline of intrahepatic transplant of Hep-53.4 cells to generate orthotopic tumours, followed by therapeutic intervention at day 14 with sorafenib (45 mg/kg), lenvatinib (10 mg/kg) or vehicle control via daily oral gavage before tumours were harvested at day 28. (B) Graph shows tumour burden of Hep-53.4 orthotopic tumours treated with vehicle control, sorafenib or lenvatinib. Data are mean ± s.e.m. from up to N=8 mice per treatment group. (C-D) Graphs show histological quantification and representative images of Hep-53.4 tumours stained for (C) Ki-67 and (D) active caspase-3 from mice treated with vehicle control, sorafenib or lenvatinib. Data are mean ± s.e.m. from up to N=8 mice per treatment group. (E) Schematic shows the timeline of PCTS/PCLS generation and subsequent treatment with vehicle control, sorafenib (10 µM – 20 µM) or lenvatinib (0.25 µM - 0.5 µM) throughout a 4-day period in a rocked Bioreactor platform. (F) Graph shows ATP production measured via CellTiter-Glo assay from PCLS and PCTS harvested at day 4 following culture with vehicle control, sorafenib (10 µM – 20 µM) or lenvatinib (0.25 µM - 0.5 µM). Data are mean ± s.e.m. from N=2 PCLS/PCTS. (G) Graph shows cytotoxicity measured via LDH assay from PCLS treated with vehicle control, sorafenib (10 µM – 20 µM) or lenvatinib (0.25 µM - 0.5 µM) across 4 days in culture. Data are mean ± s.e.m. from N=2 PCLS. (H-I) Histological quantification and representative images of (H) Ki-67-stained and (I) active capsase-3-stained PCTS following treatment with vehicle control, sorafenib (20 µM) or lenvatinib (0.5 µM) across 4 days in culture. Data are mean ± s.e.m. from up to N=4 PCTS. Scale bars: 50 μm.

To determine whether the cytostatic and apoptotic effects observed *in vivo* could be recapitulated *ex vivo*, PCLS and PCTS were generated from livers with orthotopic Hep-53.4 tumours and treated with sorafenib and lenvatinib (**Figure 3E**). Daily resazurin and CellTiter-Glo assays were performed on PCTS and PCLS to assess metabolic activity and ATP production respectively, while LDH levels were measured in media samples harvested from PCLS to examine possible cytotoxicity in non-tumour liver tissue following drug treatment. In line with the *in vivo* TKI responses, treatment of PCTS with the higher 20 μM dose of sorafenib resulted in a significant decrease in tumour slice viability characterised by a reduction in both ATP production and metabolic activity (**Figure 3F and Supplemental Figure 3B**), while analysis of LDH release demonstrated that 20 μM sorafenib was associated with hepatotoxicity in the initial 24 hours of treatment before this effect ameliorated (**Figure 3G**). Additionally, whilst no significant change was observed in cancer cell proliferation in the PCTS, an increase in active caspase-3^+^ apoptotic cells was observed following treatment with 20 μM sorafenib, reflective of the *in vivo* immunohistochemical data (**Figure 3H-I**). These data suggest that the Hep-53.4 PCTS/PCLS system can be used to identify drug-related toxicities whereby efficacious drug concentrations that are critically non-toxic can be identified.

In contrast to sorafenib, PCTS treated with lenvatinib showed no decrease in tumour cell or hepatocyte viability characterised by ATP production and metabolic activity (**Figure 3F and Supplemental Figure 3B**). Hepatotoxicity was observed in PCLS during the initial 24 hours of treatment only, before LDH release abated for the remainder of the culture period, and there was no significant impact on PCLS metabolic activity (**Figure 3G and Supplemental Figure 3C**). Mirroring the *in vivo* immunohistochemical findings, lenvatinib treatment was associated with a decrease in Ki-67^+^ proliferative cells whilst causing no change in apoptosis in PCTS (**Figure 3H-I**). Taken together these data suggest that the PCTS system largely recapitulates the TKI therapy responses observed in the *in vivo* orthotopic HCC model.

We have previously shown that Hep-53.4 orthotopic tumours are responsive to anti-PD1 immunotherapy (17). We therefore wanted to determine whether PCTS generated from orthotopic tumours would also respond to immunotherapy *ex vivo* (**Figure 4A**). Compared to IgG treated controls, anti-PD1 treated PCTS showed an increase in cell death characterised by elevated TUNEL staining (**Figure 4B**). Importantly, this effect was limited to PCTS, with anti-PD1 treated PCLS having comparable levels of TUNEL staining to IgG controls (**Figure 4B**). In line with the published *in vivo* model, there was an associated significant increase in numbers of CD3+ T cells in anti-PD1 treated PCTS (**Figure 4C**). To determine whether T cells were actively proliferating in the anti-PD1 treated PCTS, a multiplexed spatial analysis was performed. This revealed that anti-PD1 immune checkpoint blockade promotes CD8 T cell proliferation as identified by dual Ki67^+^CD8^+^ positive T cells **(Figure 4D-E)**.

**Figure 4.**
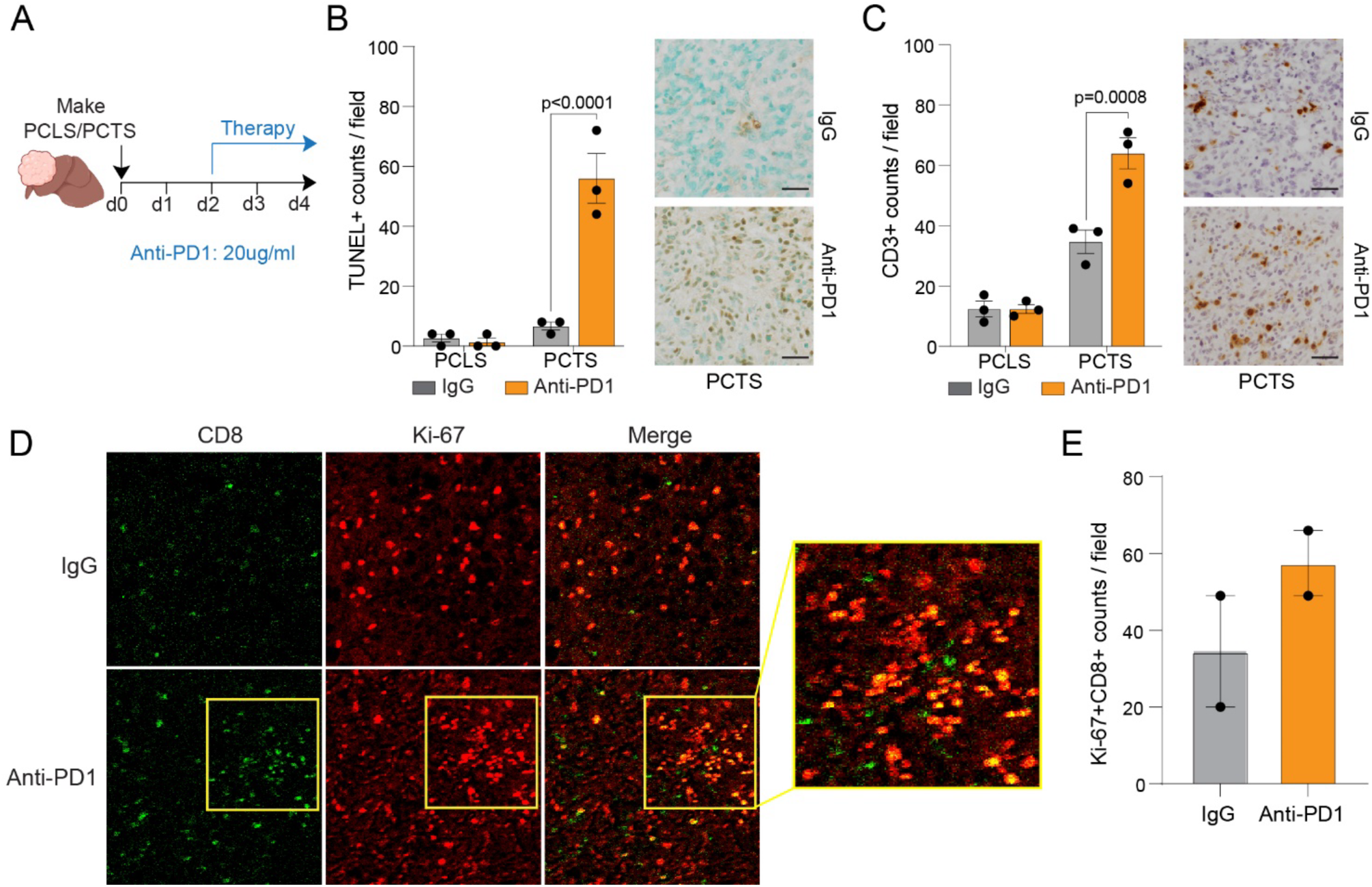
PCTS recapitulate in vivo responses to immunotherapy. (A) Schematic shows the timeline of PCTS/PCLS generation and subsequent 4-day culture period, with therapeutic intervention with anti-PD1 (20 µg/ml) from day 2 onwards. (B-C) Histological quantification and representative images of (B) TUNEL-stained and (C) CD3-stained PCLS and PCTS following culture with isotype control or anti-PD1 (20 µg/ml). Data are mean ± s.e.m. from N=3 PCLS/PCTS per treatment group. (D) Representative images from Hyperion multiplex spatial analysis of PCTS cultured with isotype control or anti-PD1 (20 µg/ml), showing CD8 (green) and Ki-67 (red). (E) Graph shows quantification of Ki-67^+^CD8^+^ T cells from Hyperion imaging mass cytometry images. Data are mean ± s.e.m. for N=2 regions of interest. Scale bars: 50 μm.

Taken together these data demonstrate that PCTS can effectively maintain the TME in culture and recapitulate both TKI and immunotherapy treatment effects observed in the *in vivo* model, therefore offering an *ex vivo* murine pre-clinical HCC platform for studying therapeutic responses.

### Modelling tumour evolution and miniaturising the PCTS platform

Given the observed robustness of *ex vivo* cultured PCTS we hypothesised that since the tumour slices remain viable for longer than slices produced from healthy mouse liver, they may not require the bioreactor system, this potentially dramatically simplifying the model and making it accessible to investigators who are unable to gain access to the bioreactor. We therefore longitudinally assessed the metabolic activity of PCTS in both the Bioreactor and standard static 12-well plates, alongside ATP production and cell death using resazurin, CellTiter-Glo and LDH assays respectively (**Figure 5A**). The data collected indicated that unlike PCLS, PCTS can be cultured in standard 12-well plates without any detrimental effect on slice viability, and with no significant differences observed in metabolic activity or ATP production between rocked and static PCTS (**Figure 5B-C**). Unexpectedly, PCTS cultured in static 12-well plates released lower levels of LDH than rocked PCTS in the Bioreactor, indicative of lower cytotoxicity which provides a significant advantage of avoiding rocking the tissue (**Figure 5D**).

**Figure 5.**
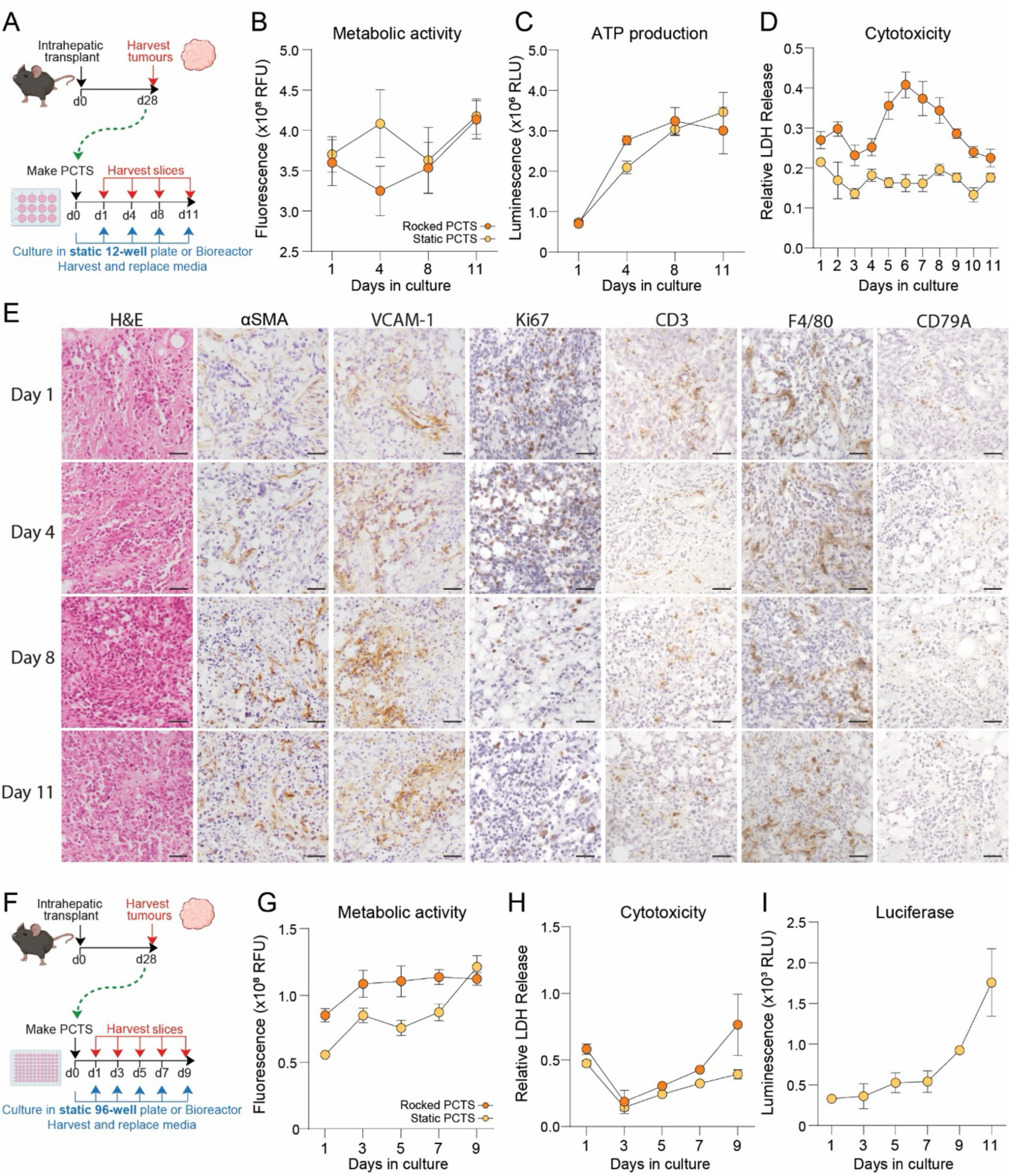
Modelling tumour evolution and miniaturisation of PCTS. (A) Schematic shows the timeline of intrahepatic transplant of Hep-53.4 cells to generate orthotopic tumours for PCTS generation, and subsequent 11-day culture period in static 12-well culture plates or the rocked Bioreactor platform. (B-D) Graphs show (B) metabolic activity, (C) ATP production and (D) cytotoxicity from PCTS cultured for 11 days in 12-well static culture or a rocked Bioreactor platform, measure via resazurin assay, CellTiter-Glo assay and LDH assay respectively. Data are mean ± s.e.m. from N=5 PCTS per group and time point. (E) Representative images of PCTS cultured in static plates at days 1, 4, 8 and 11. PCTS were H&E-stained and immunohistochemically stained for αSMA, VCAM-1, Ki-67, CD3, F4/80 and CD79A. (F) Schematic shows the timeline of intrahepatic transplant of Hep-53.4 cells to generate orthotopic tumours for the generation of miniaturised 3mm PCTS, and subsequent 9-day culture period in static 96-well culture plates or the rocked Bioreactor platform. (G-H) Graphs show (G) metabolic activity and (H) cytotoxicity from PCTS cultured for 9 days in 96-well static culture or a rocked Bioreactor platform, measured via resazurin assay and LDH assay respectively. Data are mean ± s.e.m. from N=4 PCTS per group. (I) Graph shows longitudinal luciferase assay data from SecLuc PCTS cultured in static 96-well culture plates for 11 days. Data are mean ± s.e.m. from N=4 PCTS. Scale bars: 50 μm.

To determine how the TME evolves over the extended static culture period, harvested PCTS were H&E stained and immunohistochemically stained for Ki67, αSMA, VCAM-1, CD3, F4/80 and CD79A to assess tumour cell proliferation, stromal cell activation, endothelial cell proliferation and immune cell viability (**Figure 5E**). Histological assessment of PCTS at days 1, 4, 8 and 11 revealed that TME characteristics were maintained throughout the 10-day culture period: dense tumour tissue was highlighted via H&E stain, alongside an αSMA^+^ stroma and a VCAM-1^+^ endothelium which developed throughout the culture period. Proliferative Ki67^+^ cells were still evident in the tissue at day 11, alongside CD3^+^ T cells, F4/80^+^ macrophages and CD79A^+^ B cells which were retained in PCTS harvested after 11 days.

To provide utility as a therapeutic screening platform capable of assessing a wide range of compounds, the PCTS system was miniaturised to enable medium-throughput screening of drugs. PCTS 3mm in diameter were generated as opposed to the 8mm tissue slices described previously (14), which were subsequently cultured in either static 96-well culture plates or a 96-well version of the Bioreactor for a total of 9 days (**Figure 5F**). The metabolic activity and cytotoxicity of rocked and static 3mm PCTS were determined via resazurin assay and LDH assay respectively, revealing that akin to the 12-well PCTS, 3mm PCTS do not require the rocked Bioreactor system to maintain tissue viability (**Figure 5G-H**).

We next developed a non-destructive method to allow longitudinal monitoring of PCTS viability, with the purpose of enabling dynamic drug screening. To this end, Hep-53.4 cells were transfected with a custom vector inducing the expression of secreted NanoLuc luciferase (**Supplemental Figure 1A**), which allows for longitudinal monitoring of tumour cell viability by analysing media samples that are replaced daily. Assessing media samples collected between days 1 and 11 from 3mm “SecLuc” PCTS in static culture demonstrated that increased levels of luciferase were secreted from the tissue as the culture period progressed, with an exponential increase in luciferase secretion between days 7 and 11, indicative of PCTS viability and cancer cell proliferation (**Figure 5I**).

These further innovations to the *ex vivo* PCTS model therefore provides a simplified and optimised TME culture system, whereby a greater number of therapeutic avenues can be explored using the same quantity of tumour tissue, and with a method of visualising the dynamic range of PCTS by longitudinally tracking cancer cell viability.

### Developing a medium throughput PCTS screening platform for novel drug combinations

We next determined the potential for the miniaturised SecLuc-PCTS model to assess the efficacy of a wide range of therapeutic molecules. The panel of compounds selected for the screen included molecules known to specifically target HCC, such as sorafenib, lenvatinib and anti-PD1, as well as compounds with the potential to modify other relevant pathologies such as fibrosis and steatosis in the HCC TME, and compounds capable of initiating apoptosis in cancer cells (**Figure 6A**). Miniaturised PCTS were generated from Hep-53.4 SecLuc tumours and subsequently cultured with two doses of each compound from day 1 to day 8; media was replenished daily and stored to assess luciferase levels and PCTS were harvested at day 8 to examine end-point metabolic activity and ATP levels via resazurin and CellTiter-Glo assays respectively (**Figure 6B**). Whilst many of the treated PCTS appeared viable at day 8, the first-line HCC therapy sorafenib unsurprisingly caused a drastic reduction in metabolic activity and ATP production comparable to two additional drugs in the screen, the antibiotic salinomycin and the PKC inhibitor rottlerin (**Figure 6C-D**). Comparing the ATP production and metabolic activity of PCTS in the drug screen identified a positive correlation between the two measures, with sorafenib, rottlerin and salinomycin substantially attenuating each measure of tissue viability compared to the remaining compounds (**Figure 6E**).

**Figure 6.**
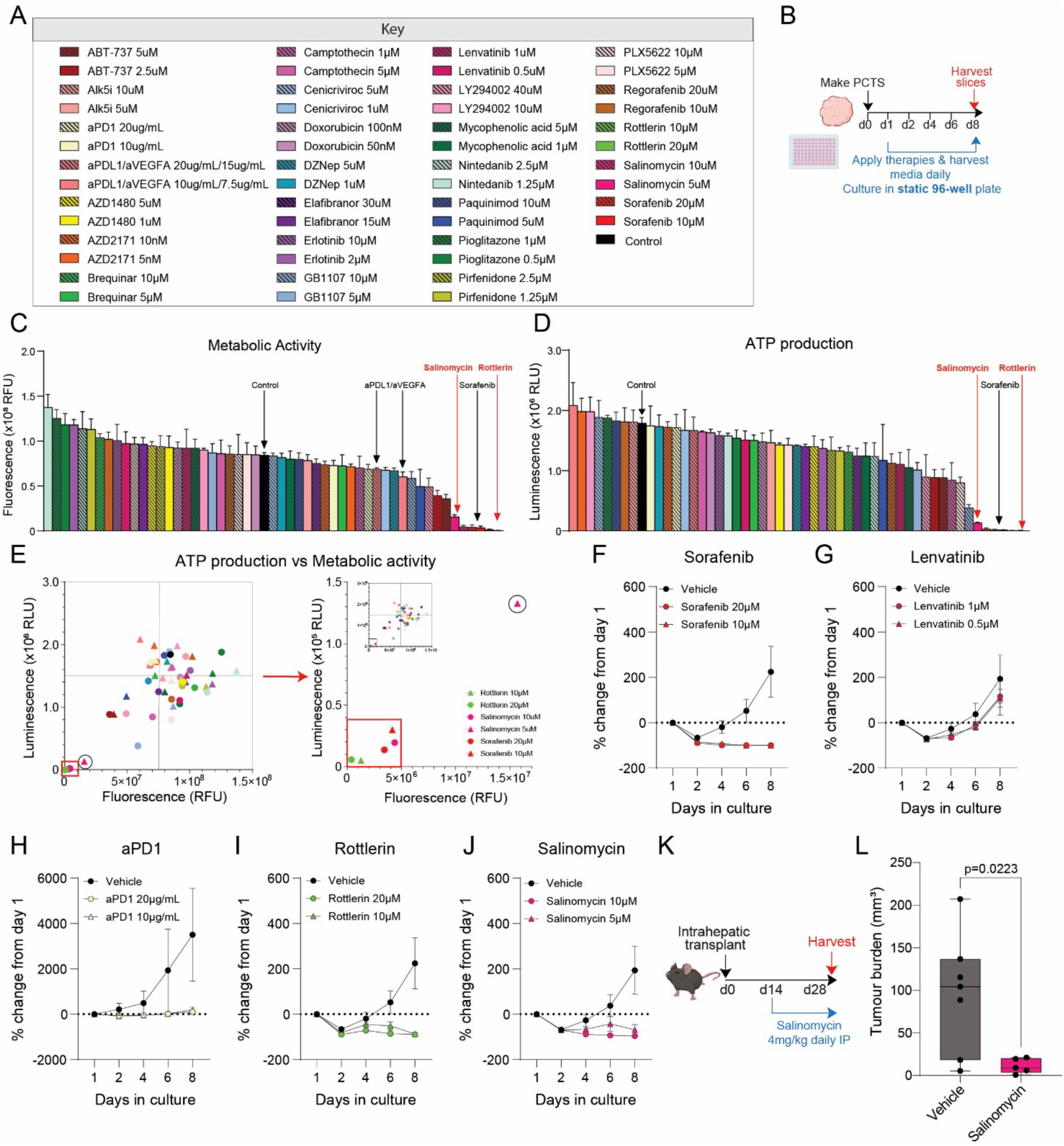
Developing a medium throughput therapeutic screening platform. (A) The key details the panel of 26 therapeutic molecules used in PCTS drug screen and the concentrations that they were applied at. (B) Schematic shows the timeline of 3mm PCTS generation and subsequent 8-day culture period in static 96-well culture plates, with 26 drugs from the screening panel applied from day 1 onwards. (C-D) Graphs show day 8 (C) metabolic activity and (D) ATP production from PCTS cultured with 26 therapies from the screening panel applied at two doses, measured via resazurin assay and CellTiter-Glo assay respectively. Data are mean ± s.e.m. for N=6 PCTS from N=5 tumours. (E) Graph shows correlation between ATP production and metabolic activity, measured by CellTiter-Glo assay and resazurin assay respectively, from PCTS cultured with 26 therapies from the screening panel applied at two doses, with emphasis on rottlerin, salinomycin and sorafenib. (F-J) Graphs show luciferase assay data from PCTS cultured with vehicle control and two doses of (F) sorafenib, (G) lenvatinib, (H) anti-PD1, (I) rottlerin and (J) salinomycin. Data are mean ± s.e.m. from N=6 PCTS from N=5 tumours. (K) Schematic shows the timeline of intrahepatic transplant of Hep-53.4 cells to generate orthotopic tumours, followed by therapeutic intervention at day 14 with salinomycin (4 mg/kg) or vehicle control via daily intraperitoneal injection, before tumours were harvested at day 28. (L) Graph shows tumour burden of Hep-53.4 tumours treated with vehicle control or salinomycin (4 mg/kg). Data are mean ± s.e.m. from up to N=7 mice per treatment group.

Implementing the secreted luciferase “SecLuc” system enabled PCTS viability to be dynamically examined throughout the time course of the therapeutic screen and avoided the use of unpaired media samples. In addition to the end-point luciferase levels, which highlighted that at day 8 most therapies resulted in lower secreted luciferase levels compared to the controls (**Supplemental Figure 4A**), the luciferase read-out provided a dynamic range illustrating that vehicle-treated PCTS secreted more luciferase as the culture period progressed from day 2 to day 8 (**Figure 6F-J**). Although the first-line TKI sorafenib completely attenuated luciferase secretion from PCTS, the first-line TKI lenvatinib failed to reduce the levels of luciferase secreted compared to the vehicle, potentially due to lenvatinib possessing greater potency against VEGF receptors alongside the lack of vasculature in PCTS (**Figure 6F-G**). Investigating PCTS response to immunotherapy demonstrated that anti-PD1 had a cytostatic effect on PCTS, and secreted luciferase levels were unchanged throughout the culture period (**Figure 6H**). Comparable to the resazurin and CellTiter-Glo data, both salinomycin and rottlerin successfully reduced the levels of secreted luciferase compared to the vehicle controls, as well as the baseline day 1 luciferase levels (**Figure 6I-J**).

The antibiotic salinomycin has previously been identified to possess anti-cancer properties, and has demonstrated efficacy in targeting murine orthotopic HepG2 tumours without reported adverse effects (22). Salinomycin was therefore selected to target Hep-53.4 tumours *in vivo*, to validate the capability of the PCTS model to identify efficacious anticancer compounds prior to conducting costly *in vivo* drug testing experiments with mice. To confirm that the cytotoxic effect of salinomycin in PCTS would not be observed in surrounding non-tumour liver tissue, PCTS and PCLS were cultured in the rocked Bioreactor system with salinomycin *ex vivo* to investigate the impact of the drug on metabolic activity (**Supplemental Figure 4B**). Resazurin assays performed on PCTS across an 8-day period illustrated that 5 μM and 10 μM salinomycin severely impacted PCTS viability by day 5 and day 7 respectively in a similar manner to 20 μM sorafenib which was used as a positive control, whilst the vehicle treated PCTS remained viable until day 8 (**Supplemental Figure 4C**). In contrast, non-tumour PCLS treated with salinomycin remained viable until day 3, comparable to vehicle treated PCLS, before all PCLS became unviable in culture at day 4 independent of salinomycin treatment (**Supplemental Figure 4D**). After determining that salinomycin did not exert obvious cytotoxic effects on non-tumour liver tissue, mice with implanted Hep-53.4 tumours were treated with 4 mg/kg salinomycin (**Figure 6K**). This resulted in a significant reduction in tumour burden compared to vehicle-control treated tumours, with macroscopic tumours just about visible following salinomycin treatment (**Figure 6L**), while there was a negligible impact on liver weight (**Supplemental Figure 4E**).

Establishing salinomycin as a drug capable of targeting *in vivo* Hep-53.4 tumours following its identification in the therapeutic screen highlights the potential of the PCTS platform to identify novel HCC therapies. This could avoid high costs, animal suffering and unnecessary use of large numbers of animals to assess compounds that do not possess potent anti-cancer activity.

## Discussion

The treatment landscape for HCC is advancing at considerable pace, evolving over the past two decades from single-agent TKIs to tailored combination therapies and the integration of immunotherapy into HCC clinics as front-line systemic therapies. With a similar pace of advance with pre-clinical studies and new insights into the biology and immunology of HCC, we can expect new therapeutic targets to continue to emerge and inform the design of novel drugs, biologics, and combination strategies. Experimental mouse models of HCC are currently the mainstay preclinical platform for determining the likely efficacy of novel drugs and combination therapies. While such models can be a powerful translational tool, they have inherent limitations, not least their high cost and the incurred suffering of very high numbers of animals. In addition, *in vivo* mouse models of most cancers do not readily lend themselves to ‘at-scale’ drug screens or for the robust testing of experimental combinatorial therapies where determining the effective dosage and timing of administration of the individual drugs can be critical in achieving an optimal therapeutic outcome (23). With HCC as our paradigm disease, we have produced a viable and accessible solution to the limitations of *in vivo* oncology studies in mice by optimising the conditions for use of *ex-vivo* live PCTS including their miniaturisation to 96-well culture scale and development of dynamic quantification of tumour cell viability.

PCTS are produced by precision cutting of thin slices of *ex-vivo* tumour tissues using a vibrating microtome and have previously been reported for to maintain the architecture and diverse cellular constituents of the TME (24). PCTS have also been reported to maintain T cell populations (CD4^+^ and CD8+) of the original tumour and to have the potential for immunotherapy investigations (24). Both human and mouse PCTS have been described for a variety of tumours, however their application for drug screening is yet to be fully realised. Concerning HCC, Jagatia and colleagues recently described the design of human PCTS as a platform for preclinical drug testing in primary liver cancer samples (15). This latter study described maintenance of HCC- and cholangiocarcinoma-derived PCTS for up to 8 days in culture with retention of proliferating tumour cells, stromal cells and continued presence of intratumoral CD3^+^ and CD45^+^ leukocytes. The authors also validated HCC-PCTS as a model for pre-clinical testing of TKIs and anti-PD1. However, despite this advance there are considerable limitations that impact on the wider utility of human PCTS, not least the availability of fresh human tumour tissue which requires close location of a surgical oncology unit and logistic solutions for the timely acquisition of the tissues from operating theatres, and the requirement for clinical pathology assessment of tissues prior to their release for transfer to the laboratory. Furthermore, any pre-clinical oncology model that is to be utilised for drug development purposes requires standardisation to overcome inherent variables that influence the biology of the PCTS and their responses to drug treatments, such as intra- and inter-tumour heterogeneity (both architecturally and in terms of cellular and stromal tissue constituents). These variables are extremely challenging to overcome for many human solid tumours including HCC. We therefore explored a rationale alternative of combining the standardisation and ease of accessibility of mouse modelling with optimisation and scaling of *ex-vivo* mouse HCC-PCTS to produce a 3-dimensional TME platform that can be used in most laboratories, including in an industry setting where access to human tumour tissue can be rare and unpredictable.

As with human HCC-PCTS, tumour slices from the orthotopic HCCs retained proliferative epithelial tumour cells, stromal cells plus a variety of immune cells including cytotoxic CD8^+^ T cells and tumour-associated macrophages throughout an extended culture period of at least 11 days. This 11-day culture period provides a practical time window for extensive testing of small molecule oncology drugs and immunotherapy agents including strategizing the sequencing of dosing regimens in the context of combinatorial approaches. With respect to immune checkpoint blockade we were able to demonstrate not only an anti-tumour response to anti-PD1, but also showed that resident CD8^+^ T cells expanded in numbers and to include Ki67^+^ cells indicative of induced proliferation occurring within the cultured TME. As we have previously described the requirement of a rocked bioreactor system for optimal maintenance of rat and human liver slices (14) we initially cultured HCC-PCTS under these conditions, however subsequent experiments indicated no advantage of the bioreactor or from the prevention of contact of tumour slices with a plastic substrate. Given the ease of optimisation of HCC-PCTS under standard cell culture conditions we were able to make adaptions on the platform to enable drug screening in the context of an intact live TME. To allow for dynamic monitoring of effects of drugs on tumour cell numbers we generated Hep-53.4 tumour cells expressing secreted luciferase and employed these cells for producing orthotopic tumours. As PCTS media is changed daily the growth of tumour cells is then easily determined across a time course, and as we were able to demonstrate, can be used to monitor the kinetics of TKI cytotoxic effects by measurement of luciferase activity in the culture media. The second modification was to miniaturise the platform to 96-well scale, this enabled us to screen 26 therapeutic agents at two doses and with inclusion of appropriate vehicle or antibody controls. The screen was carried in replicates requiring a total of 5 *ex vivo* tumours. A similar screen carried out *in vivo* would require 520 test mice in addition to vehicle and antibody controls (20 mice), when using 10 mice per group, this highlighting the potential for *ex vivo* PCTS to reduce animal usage for oncology as well as enabling at-scale drug screening in an intact mouse TME. The screen successfully identified two drugs (rottlerin and salinomycin) that are not currently used in the HCC clinic, one of the drugs, salinomycin, was then validated as effective for reducing tumour burden *in vivo*. These PCTS adaptions therefore provide the opportunity for real-time monitoring of drug screens at medium-throughput scale and most importantly in the context of a live and anatomically faithful TME that maintains the key cell types that are targeted by small molecule drugs and immunotherapy agents. Of note, PCTS can be processed downstream of a drug study for flow cytometry, live cell imaging, generating single cell transcriptomics data, and producing spatial omics data which if combined will exploit the intact live TME architecture of tumour slices to report effects of therapeutic molecules with a more wholistic approach than can possibly be achieved with simpler 2D or 3D cell culture systems including organoids. It will also be possible to genetically modify the Hep53.4 cell line, for example by CRISPR/Cas9 technology to allow drugs to be screened in the context of mutations commonly found in human HCC tumours. Such an approach can improve our depth of understanding of the mechanisms by which oncology agents bring about their beneficial effects but also predict toxicities and unwanted influences on the viability and phenotypes of tumour-associated immune cells. These insights can then be employed to improve the design of future cancer therapeutics.

In summary, we report a dynamic 96-well scale murine HCC TME platform for oncology biology and drug development purposes that can be readily established and employed by the majority of academic and industry laboratories that have access to an animal unit, appropriate ethical and regulatory approvals and standard cell culture facilities. As we were able to routinely generate a full 96-well plate of PCTS from a single tumour, the platform delivers an estimated 98.7-fold reduction (assuming n=4 PCTS per group for 11 compounds & 1 vehicle control) in use of animals as well as avoiding exposing mice to suffering caused by the toxic effects of experimental drugs and novel combination treatments. We envisage the platform integrating downstream of higher throughput human tumour cell culture systems such as patient-derived organoids to validate drug efficacies and explore mechanism of drugs in the intact TME, this replacing the need for *in vivo* pre-clinical studies as part of the drug development process. The adaptions we report could also be applied to other primary as well as secondary tumours of the liver. Moreover, with advancing improvements in the engineering of humanised mice and their combination with implanted patient tumour tissues, the PCTS platform can be further developed to deliver personalised oncology at a scale that allows for examination of complex combinatorial drug dosing and sequencing.

## Abbreviations

ALD: Alcohol-related liver disease
AST: Aspartate aminotransferase
H&E: haematoxylin and eosin
HCC: hepatocellular carcinoma
IMC: Imaging Mass Cytometry
LDH: lactate dehydrogenase
MASLD: metabolic dysfunction-associated steatotic liver disease
MASH: metabolic associated steatohepatitis
PCLS: Precision cut liver slices
PCTS: precision cut tumour slices
RLTD: relative level of transcriptional difference
TUNEL: Terminal deoxynucleotidyl transferase-mediated dUTP nick end
TME: tumour microenvironment
TKIs: tyrosine kinase inhibitors
WT: wild type

## Author contributions

JL, FO and DAM contributed equally to the experimental design, data interpretation and writing of the manuscript. JL and AC led the laboratory work and AC together with JL and FO prepared the figures. AC contributed to the writing of the manuscript. KK carried out a large volume of the experimental work and was assisted by ER-G and AF with Hyperion IMC. ET, DG, LAB, HP, RC, SL, EK and DS made important technical contributions to the work that were crucial to the study and worthy of co-authorship. All authors read and approved the final manuscript.

## Acknowledgements

We would like to thank the Newcastle University Comparative Biology Centre and Flow Cytometry Core Facility for their technical assistance.

---

**SF1.**
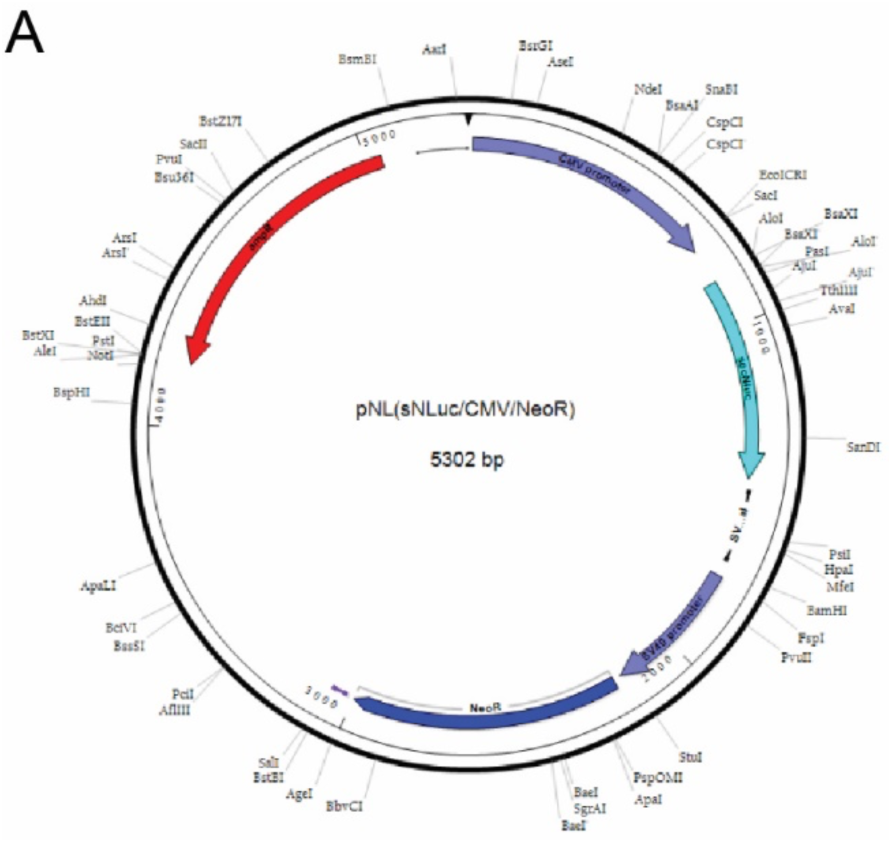
(A) Vector map of custom plasmid (Promega) inducing the expression of secreted NanoLuc luciferase.

**SF2.**
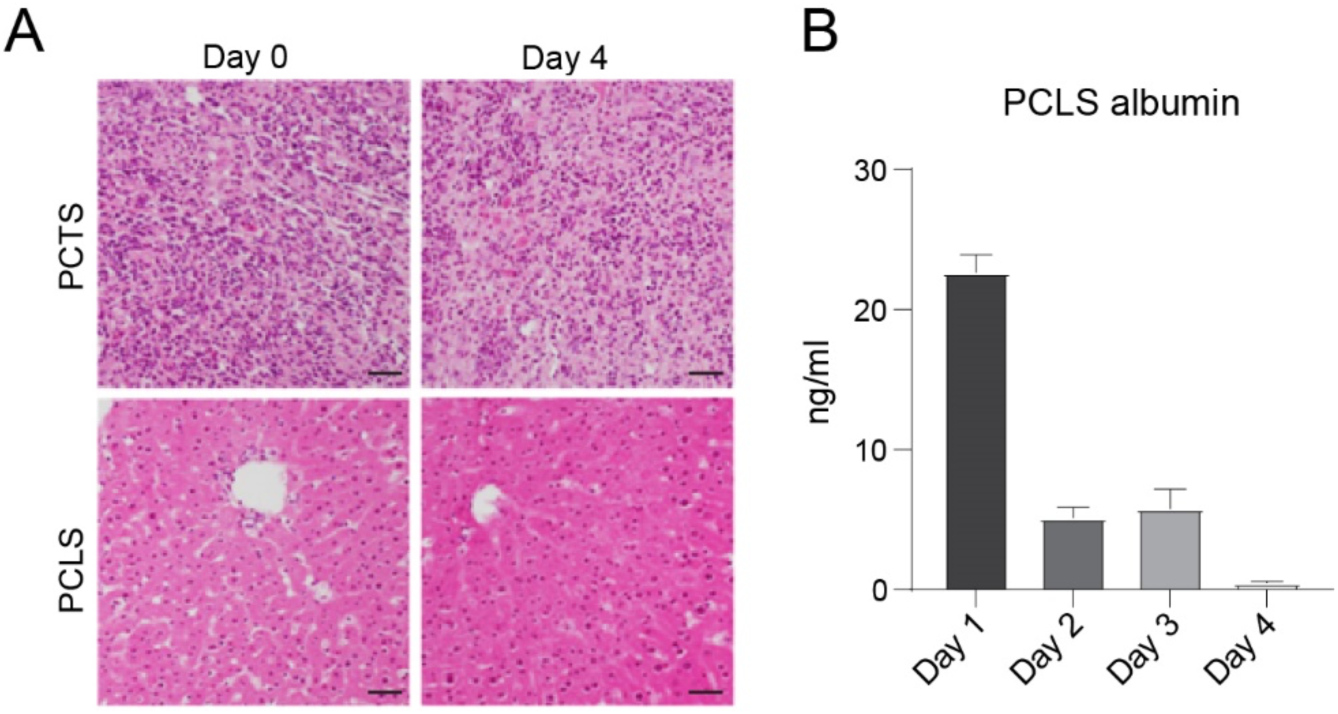
(A) Representative images of H&E-stained PCTS and PCLS at day 0 and day 4 following culture in a rocked Bioreactor platform. (B) Graph shows concentration of soluble albumin secreted from PCLS cultured for 4 days in the rocked Bioreactor platform. Data are mean ± s.e.m. from up to N=20 paired PCLS. Scale bars: 50 μm.

**SF3.**
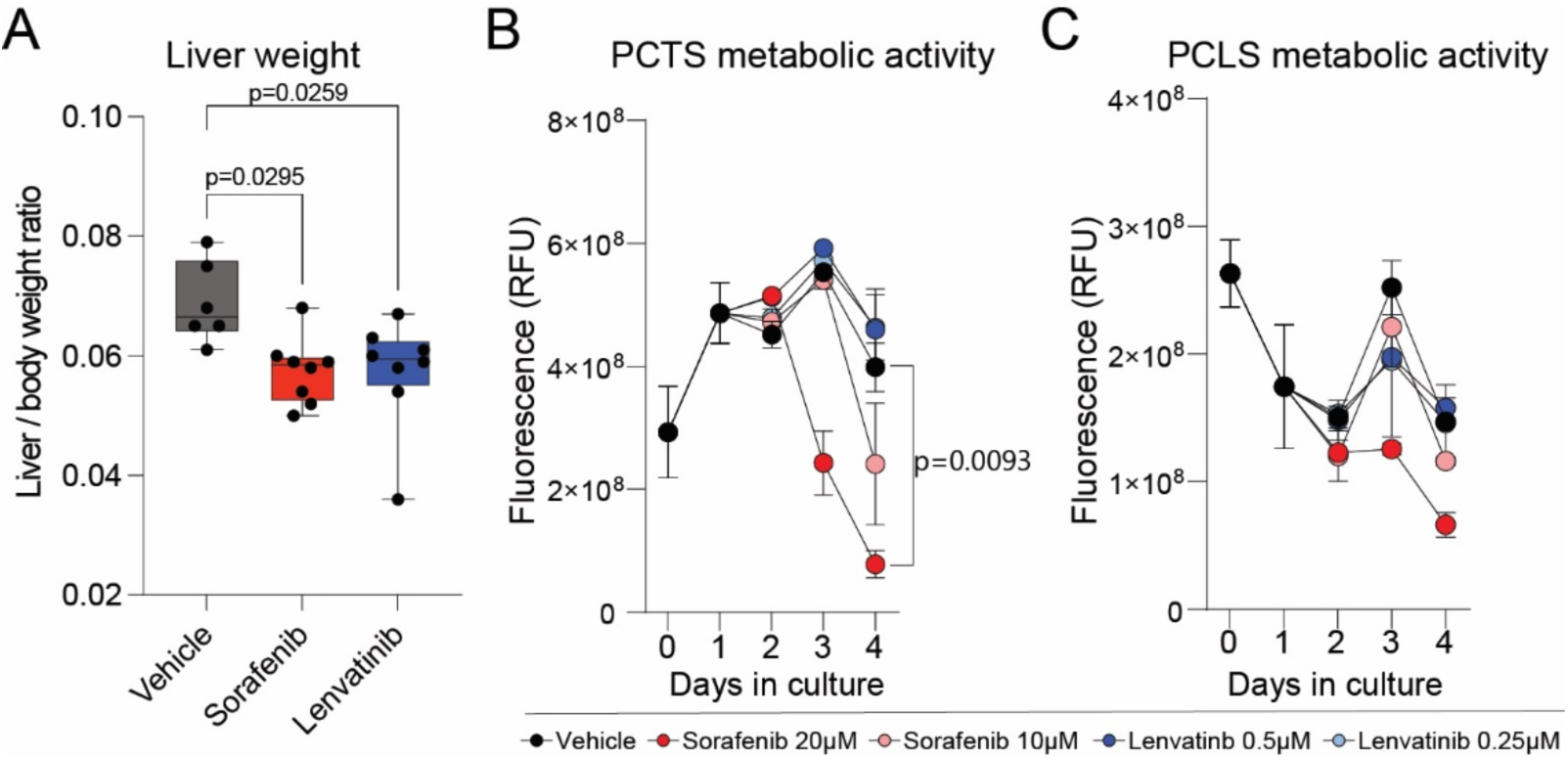
(A) Graph shows liver to body weight ratio of mice with Hep-53.4 orthotopic tumours following treatment with vehicle control, sorafenib (45 mg/kg) or lenvatinib (10 mg/kg). Data are mean ± s.e.m. from up to N=8 mice per treatment group. (B-C) Graphs show metabolic activity of (B) PCTS and (C) PCLS cultured for 4 days with vehicle control, sorafenib (10 µM – 20 µM) or lenvatinib (0.25 µM - 0.5 µM), measured via resazurin assay. Data are mean ± s.e.m. from up to up to N=3 PCTS/PCLS per treatment group and time point.

**SF4.**
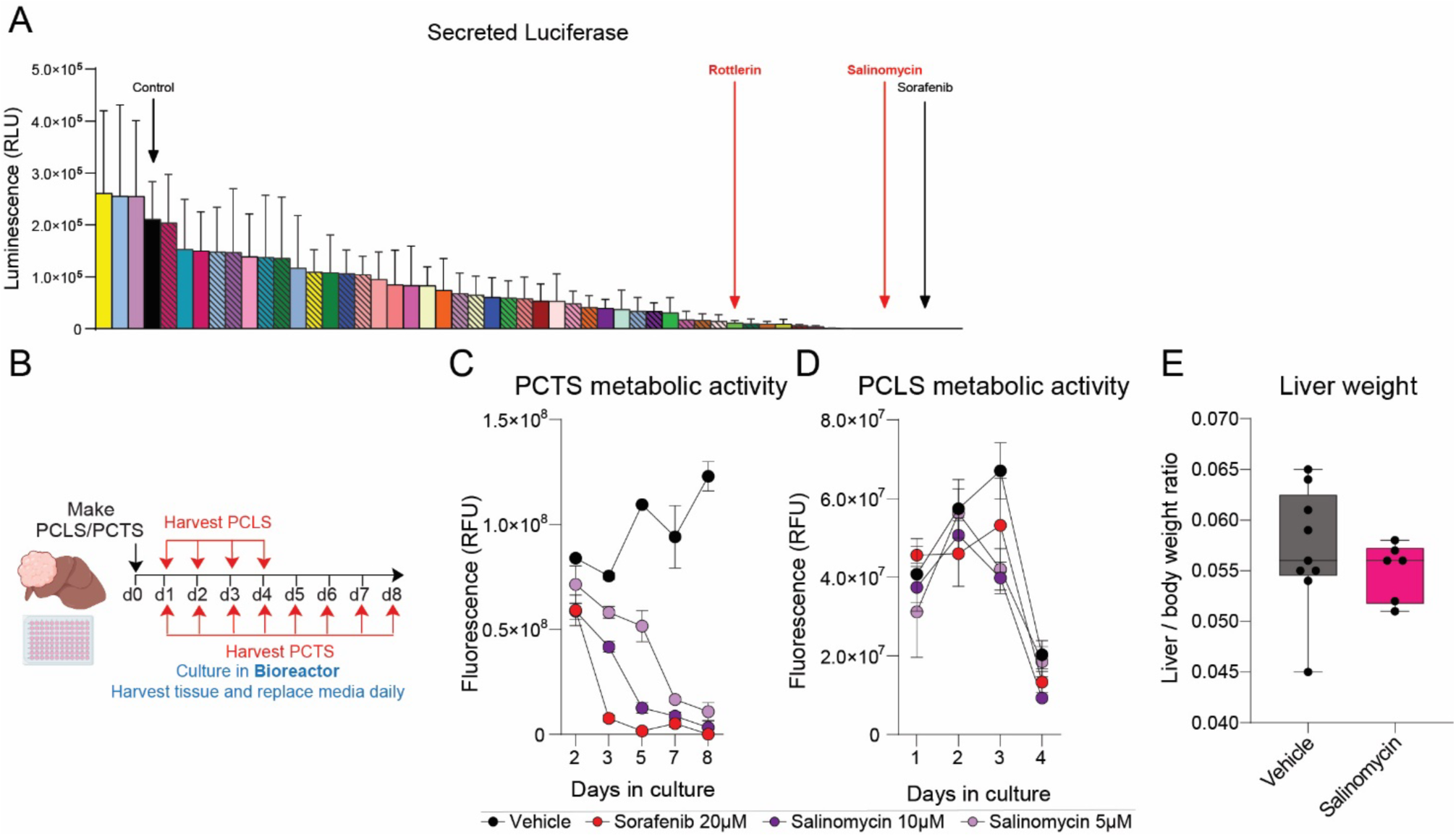
(A) Graph shows day 8 secreted luciferase levels from PCTS cultured with 26 therapies from the screening panel applied at two doses, measured via Nano-Glo luciferase assay. Data are mean ± s.e.m. for N=6 PCTS from N=5 tumours. (B) Schematic shows the timeline of PCLS and PCTS generation and subsequent culture period of 4 and 8 days respectively. (C-D) Graphs show metabolic activity of (C) PCTS and (D) PCLS cultured for 8 and 4 days respectively with vehicle control, sorafenib (20 µM) or salinomycin (5 µM – 10 µM), measured via resazurin assay. Data are mean ± s.e.m. from up to N=4 PCTS/PCLS per treatment group and time point. (E) Graph shows liver to body weight ratio of mice with Hep-53.4 orthotopic tumours following treatment with vehicle control or salinomycin (4 mg/kg). Data are mean ± s.e.m. from up to N=8 mice per treatment group.

